# Preclinical and Toxicology Assessment of ISFP10, an Inhibitor of Fungal Phosphoglucomutase (PGM)

**DOI:** 10.1101/2025.01.07.631740

**Authors:** Madeline R. Pellegrino, Marc Bindzus, Theodore J. Kottom, Savita Ayyalasomayajula, Eunhee S. Yi, Andrew H. Limper

## Abstract

**Background and Objective:** Previously, the novel small molecule ISFP10 has been shown to inhibit fungal phosphoglucomutase (PGM) activity in *Aspergillus fumigatus* and *Pneumocystis* spp. With 50-fold selectivity over the human PGM molecule due to the presence of a unique yet conserved cysteine residue present in a number pathogenic fungal PGMs, use of this compound may provide a novel broad-spectrum approach to treating fungal infections. Accordingly, we sought to determine the tolerability in test animals receiving this compound, as well as the potential antifungal activity of ISFP10 on cultures of the common fungal pathogens *Candida albicans* and *Candida glabrata*.

**Methods:** C57BL6 mice received once daily intraperitoneal (IP) injections of 100 mL of vehicle control (DMSO) or ISFP10 at a concentration of 10 mg/kg. Body weights were recorded daily for 7 days of treatment. On the final day, mice were weighed and euthanized. Postmortem blood collection was conducted via cardiac puncture and distributed to EDTA and lithium heparin tubes for complete blood count (CBC) and comprehensive blood chemistry panels, respectively. Liver, kidney, and lung tissue were also harvested and placed in 10% formalin for H&E staining and blinded histopathologic scoring. Lung samples were further analyzed for proinflammatory cytokines using enzyme-linked immunosorbent assays (ELISA) and quantitative PCR (qPCR). Furthermore, ISPF10 was tested for antifungal activity via 8-hour growth curve analysis in a concentration-dependent fashion against *Candida albicans* and *Candida glabrata*.

**Results:** There was no significant difference in the daily or final body weights of the mice receiving 10 mg/kg of ISFP10 compared to those of the vehicle control group. Extracellular matrix (ECM) transcripts for IL-6 and TNFα were statistically similar via qPCR. ELISA results of proinflammatory cytokines for IL-6 was not significant whereas TNFα levels in lung tissue from the ISFP10 treatment group were significantly reduced, indicating a potential anti-inflammatory effect of ISFP10 at this dosage. Overall, blood chemistry and CBC analysis revealed no overall significant differences between the two groups, except for increased neutrophil counts and decreased potassium levels in samples collected from ISFP10 treated animals compared to the vehicle control group. These laboratory abnormalities were not of clinical significance to the test animals. Blinded histopathological examination revealed no abnormalities or evidence of critical organ toxicity from all groups. Inhibition of *C. albicans* and *C. glabrata* culture growth by ISFP10 was concentration-dependent in YPD liquid media containing the ISFP10 compared to vehicle control.

**Conclusions:** Our preliminary testing of ISFP10 revealed no inherent safety or toxicology concerns within the observed parameters. These data further support significant culture suppressive activity against *C. albicans and C. glabrata*. Taken together, these observations of ISFP10 further indicate that targeting PGM might be a novel and viable therapeutic strategy for serious fungal infections.

**Key points:** An inhibitor specific to fungal PGM enzymes, termed ISFP10, was generally well tolerated when administered via intraperitoneal (IP) injection in mice.

ISFP10 displays antifungal activity in a concentration-dependent manner against *C. albicans* and *C. glabrata*.

## 1. Introduction

Fungal cell wall generation is an appealing target for antifungal therapy due to its absence in mammals [1]. Many medically relevant fungi, contain a glucan-rich cell wall which is synthesized from uridine diphosphate glucose (UDP), requiring the conversion of glucose-6-phosphate to glucose-1-phosphate by the enzyme phosphoglucomutase (PGM)[2]. Recently, it has been shown that in *A. fumigatus*, PGM is essential for organism viability and cell wall integrity, and that selective targeting of a cysteine residue of the *A. fumigatus* PGM (*Af*PGM) with a compound termed ISFP10 can inhibit *Af*PGM activity with an IC_50_ of 2 μM and with at least 50-fold selectivity over the human homolog (which lacks the conserved cysteine)[3]. We have previously published that *P. jirovecii* and *P. murina* also possess PGM enzymes with this conserved cysteine [2]. These data provide initial proof of concept that selective targeting PGM in *Pneumocystis* versus the mammalian homolog, might be a novel therapeutic strategy to treat Pneumocystis pneumonia (PCP). This study sought to evaluate host responses to the short-term administration of ISFP10 in mice via IP injection, using toxicological, inflammatory, and tissue analyses to determine overall tolerability of the agent. Finally, we demonstrate that ISFP10 can reduce the growth of *C. albicans* and *C. glabrata*, two common fungal pathogens *in vitro*. The data presented here demonstrate the safety of ISFP10 in mice and supports the hypothesis of selectively targeting fungal PGM using *in vivo* mouse models of fungal infection for preclinical development of this novel therapeutic class.

## 2. Methods

### 2.1. Animals

10 male and 10 female 12-week-old C57BL6 mice from the Jackson Lab were used in the toxicology study. All animal procedures were performed in accordance with the Laboratory Animal Welfare Act, the Guide for the Care and Use of Laboratory Animals Welfare Act, and the Mayo Clinic Institutional Animal Care and Use Committee (IACUC) (Approval number: A00007622-24). The mice were separated by gender and housed with two to five animals per cage. After arriving at the animal housing facility, mice underwent a mandatory 2-day acclimation period before any experimentation was performed. The mice were fed PicoLab^®^ Mouse Diet 20 5058* *ad libitum* and sterile water was provided via Hydropac pouches. Each cage contained “standard enrichment material” that consisted of a Twist-n’Rich™ and a Bed-r’Nest^®^ (The Andersons Plant Nutrient Group).

### 2.2. Administration of ISFP10

ISFP10 was synthesized from SYNthesis Med Chem, Inc. (Hong Kong). Due to the lack of solubility of the inhibitor in water or saline, the inhibitor was prepared first by dissolving the compound in 100% dimethyl sulfoxide (DMSO), followed by the addition of 0.5% Methocel in 0.9% NaCl for injection at a 10%/90% vol/vol ratio. Intraperitoneal (IP) treatment (100 μl) with the DMSO/Methocel (vehicle, control mice group) or the 20 mg/kg of ISFP10 inhibitor in DMSO/Methocel was initiated on day 0 and subsequentially every day for 7 consecutive days. At day 8, mice were sacrificed, and subsequent analysis performed as described.

### 2.3. Cytokine levels determined by ELISA

Analysis of cytokine release of TNFα and IL-6 from lung protein lysates was performed using enzyme-linked immunosorbent assay (ELISA) kits purchased from Thermo Fisher Scientific [4].

### 2.4. Quantitative polymerase chain reaction analysis

For RNA extraction, tissues were lysed and homogenized in Buffer RLT Plus, (supplied with the RNeasy^®^ Plus Mini Kit; Qiagen). The lysate was then centrifuged through a genomic DNA eliminator column and 70% ethanol was added to the flow-through. The flow-through was then added to a RNeasy spin column and eluted RNA was isolated according to the manufacturer instructions. An iScript™ Select cDNA synthesis kit (Bio-Rad, Hercules, CA) that employs oligo (dT primers) and random hexamer primer mix was then used for reverse transcription. The quantitative real-time PCR employed a SYBR green PCR kit (Bio-Rad) and was performed and analyzed on a CFX96™ Real-Time PCR Detection System (Bio-Rad) [4]. Primer pairs sequences are provided in Supplementary Table 1.

### 2.5. Biochemical analysis

Following animal euthanasia, a comprehensive metabolic panel and CBC were performed on whole blood within an hour of collection via cardiac puncture. The Piccolo Xpress™ Chemistry Analyzer was utilized according to manufacturer instructions [4] along with the Abaxis VetScan HM5 Analyzer for veterinary hematology [5].

### 2.6. Histology analysis

Histopathological analysis was performed on whole kidney, liver, and lung samples fixed in 10% neutral formalin. Organs were processed in the Mayo Clinic Histology Core in Scottsdale, AZ for paraffin embedding, sectioning, and hematoxylin and eosin (H&E) staining of 5 µm sections. Blinded slide evaluations were performed by a Mayo Pathologist (ESY) to visually assess for inflammatory cell infiltration and tissue damage in critical organs. The whole organ sections presented on the slide section were graded based on the following scores: 4+, heavy aggregates and alveolar exudate; 3+, mild alveolar aggregates; 2+, heavy perivascular aggregates; 1+, mild perivascular aggregates; and 0, normal [4, 6].

### 2.7. Candida spp. growth assays

To determine the effect of ISFP10 on fungal growth, we treated two *Candida* strains with increasing concentrations of the ISFP10 inhibitor and analyzed fungal growth by absorbance at OD600_nm_ over an 8-hour time course. *C. albicans* (ATCC 90028) and *C. glabrata* (BG1182) were grown overnight in liquid YPD media at 30^°^C with continuous shaking. The cultures were then standardized to an OD600_nm_ of 0.3 before the addition of ISFP10 to achieve final concentrations of 200µM, 100µM, and 50µM. DMSO was used as a vehicle control with a concentration equivalent to the highest experimental dose. One milliliter of culture was placed in a cuvette and measured at OD600_nm_ hourly on the SpectraMax M2e microplate reader from Molecular Devices. The aliquot was returned to the incubator in between measurements. Three to four experimental runs were recorded over multiple days, and the hourly means were calculated to create a growth curve for each strain.

### 2.8. Statistical Analysis

Multigroup data were initially analyzed using analysis of variance (ANOVA) to identify overall differences. If significant differences were determined by ANOVA, additional group comparisons were conducted using a two-sample unpaired Student *t*-test for normally distributed variables. Each group (n=10) included equal numbers of 10 male and 10 female mice. Samples from each individual mouse were utilized for ELISA and qPCR analysis, and blood from 8 mice per group was analyzed for CBC and blood chemistry values.

## 3. Results

### 3.1. Administration of ISFP10 Injections Resulted in No Significant Weight Loss

There was no significant change in mouse weight observed over the 7 consecutive days of administering ISFP10 at 10 mg/kg intraperitoneally (**Fig 1**). In addition, the mice appeared vigorous and displayed normal activity during the treatment period.

**Fig 1.**
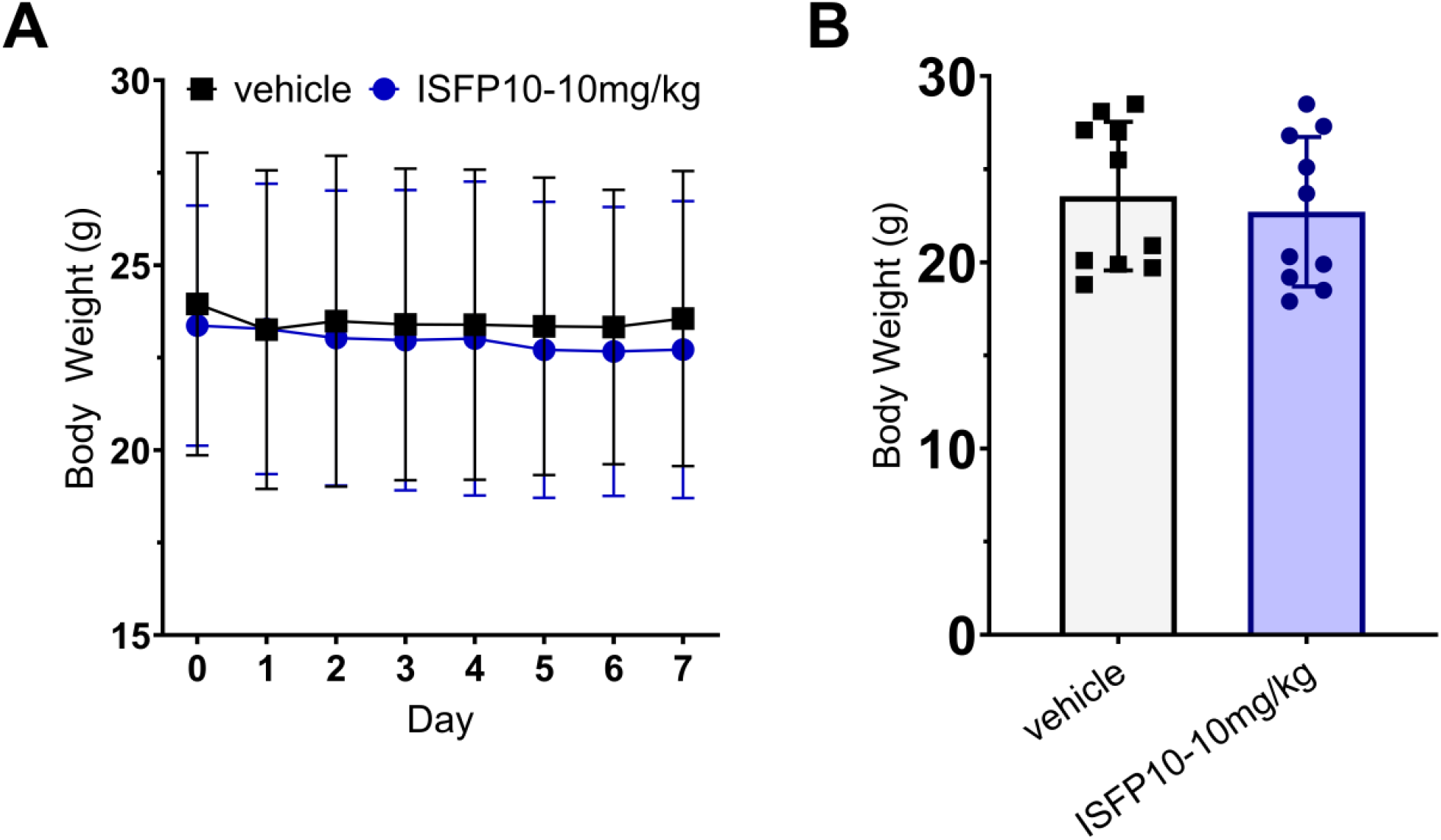
Mouse body weight with IP administration daily of ISFP10. (A) Depicts the daily weights of the vehicle control versus the 10 mg/kg doses of IP administered ISFP10 daily for 7 days. (B) Shows the final weights for the 2 groups after 8 days. (n = 10 mice/group). No significant weight changes were noted between the two groups.

### 3.2. Measurement of Lung Cytokines in IP Administration of ISFP10 vs Control Groups

To further exclude any tissue injury and inflammation induced by ISFP10, we also analyzed inflammatory cytokine levels in lung tissue. Lung was chosen as a test tissue, due to our interest in using the agent in future studies of lung infection. In whole lung lysates, the production of the inflammatory cytokines IL-6 and TNFα was measured as shown in (**Fig. 2**A-B). For IL-6, no significant difference was observed from the control group. For TNFα, there was a significant decrease in production of this cytokine with administration of ISFP10. The decrease in TNFα was not expected and might indicate an unexpected anti-inflammatory activity of ISFP10.

**Fig 2.**
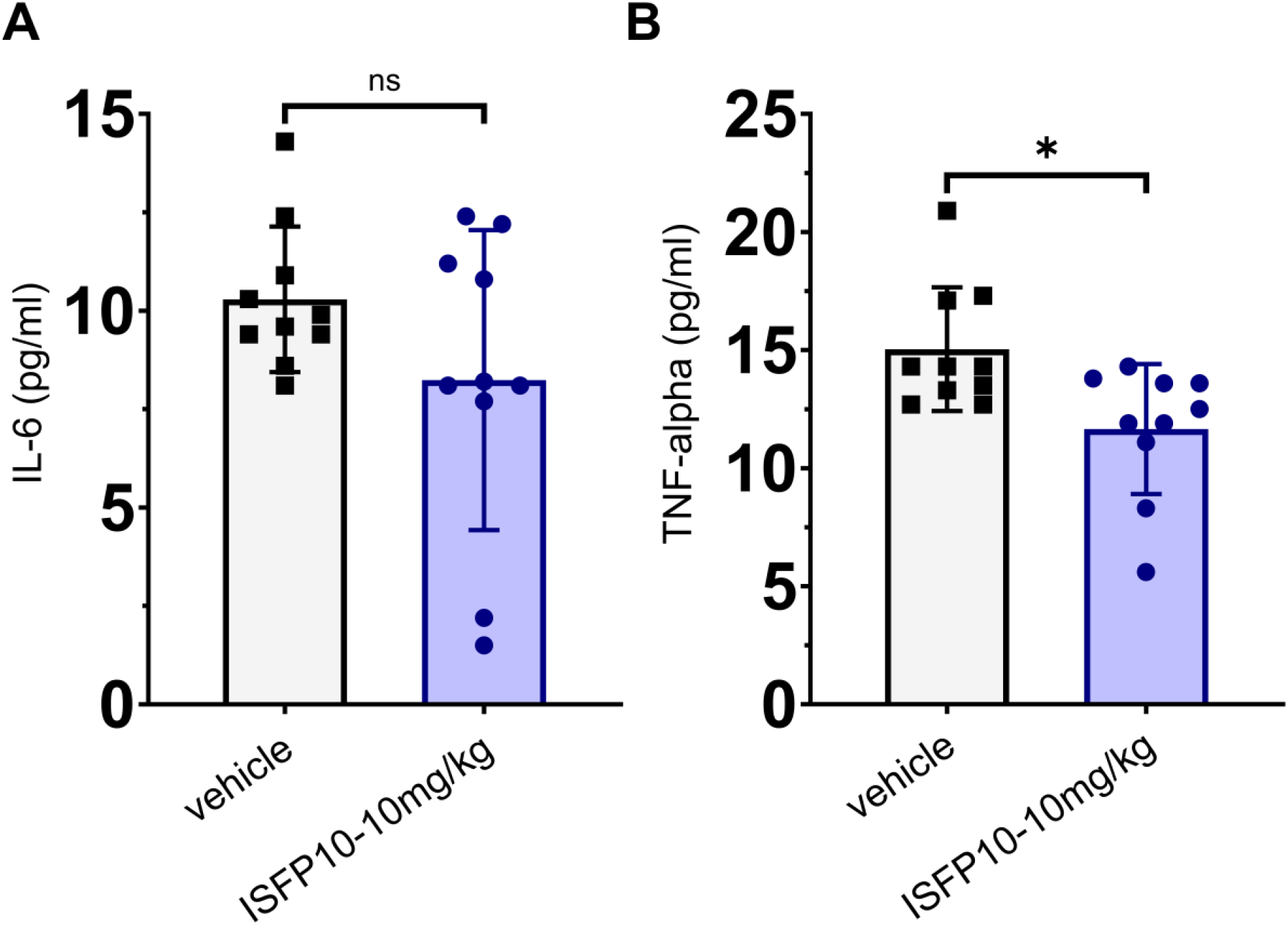
Lung proinflammatory cytokine determination post ISFP10 administration. (A) IL-6 and (B) TNFα measurements were measured by ELISA on total lung protein lysates from day 8 of the experiment (n=10 mice/group). No significant differences were seen for IL-6 production, while TNFα production was significantly reduced in the ISFP10 treatment group. ± SEM; \**p* < 0.05.

### 3.3. mRNA Analysis of Extracellular Matrix Generation (ECM)

To further exclude any subtle injuries involving fibrotic repair of the lung, we evaluated extracellular matrix gene expression in lung tissue. The mRNA expression of Fibronectin (*Fn*) and Collagen Type Alpha 1 Chain (*Col1a1*) were both determined using qPCR, with Beta-2 Macroglobulin (*B2M*) serving as the housekeeping control gene. These genes are markers of profibrotic activity as their related proteins are deposited in the ECM [7]. The results showed no significant alterations in the ECM composition of lung tissue from either group as shown in (**Fig. 3** A-B).

**Fig 3.**
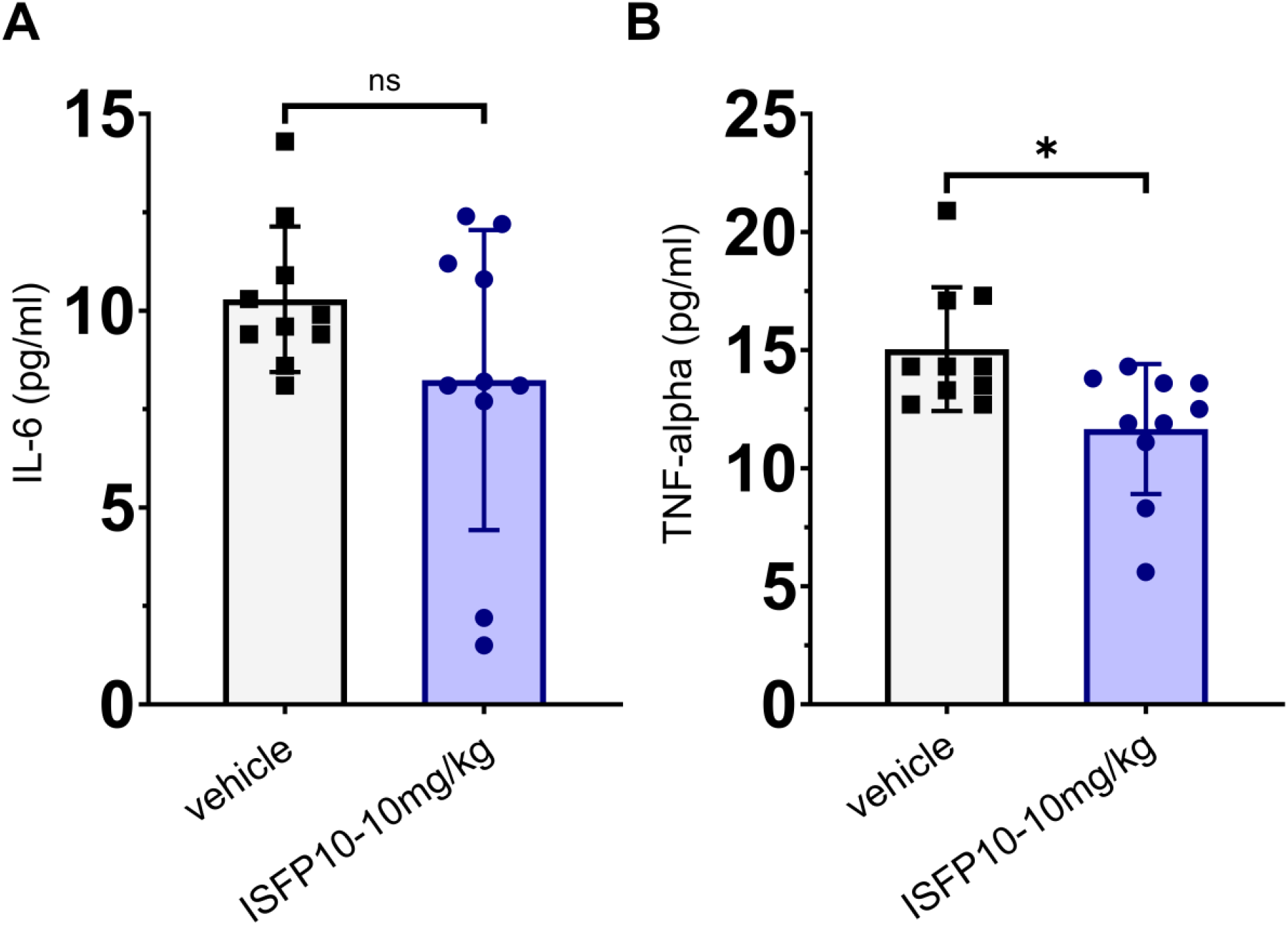
*Col1a1* and *Fn* mRNA quantification in total lung RNA post vehicle and ISFP10 administration for 7 days. Expression of (A) *Col1a1* and (B) *Fn* relative to *B2M* in total lung RNA (n=10 mice/group). No significant changes were noted with ISFP10 administration.

### 3.4. Complete Blood Count and Comprehensive Metabolic Panel Analysis

The analysis of blood collected from each treatment group revealed no significant differences in any parameter except neutrophil count and potassium ion levels (**Fig. 4 and 5**) The mice receiving IP administration of ISFP10 had significantly increased neutrophil counts (**Fig. 5** D); however, their results were not outside of the normal physiological range for the strain [8]. The mechanisms that underlying the neutrophil increase are unknow but may relate to increased mobilization of leukocytes from the bone marrow or changes in vascular distribution of the cell population. Conversely, the chemistry panel showed significantly reduced potassium ion values in the ISPF10 group (**Fig. 4** B) but once again, not also to any degree of clinical significance [8].

**Fig 4.**
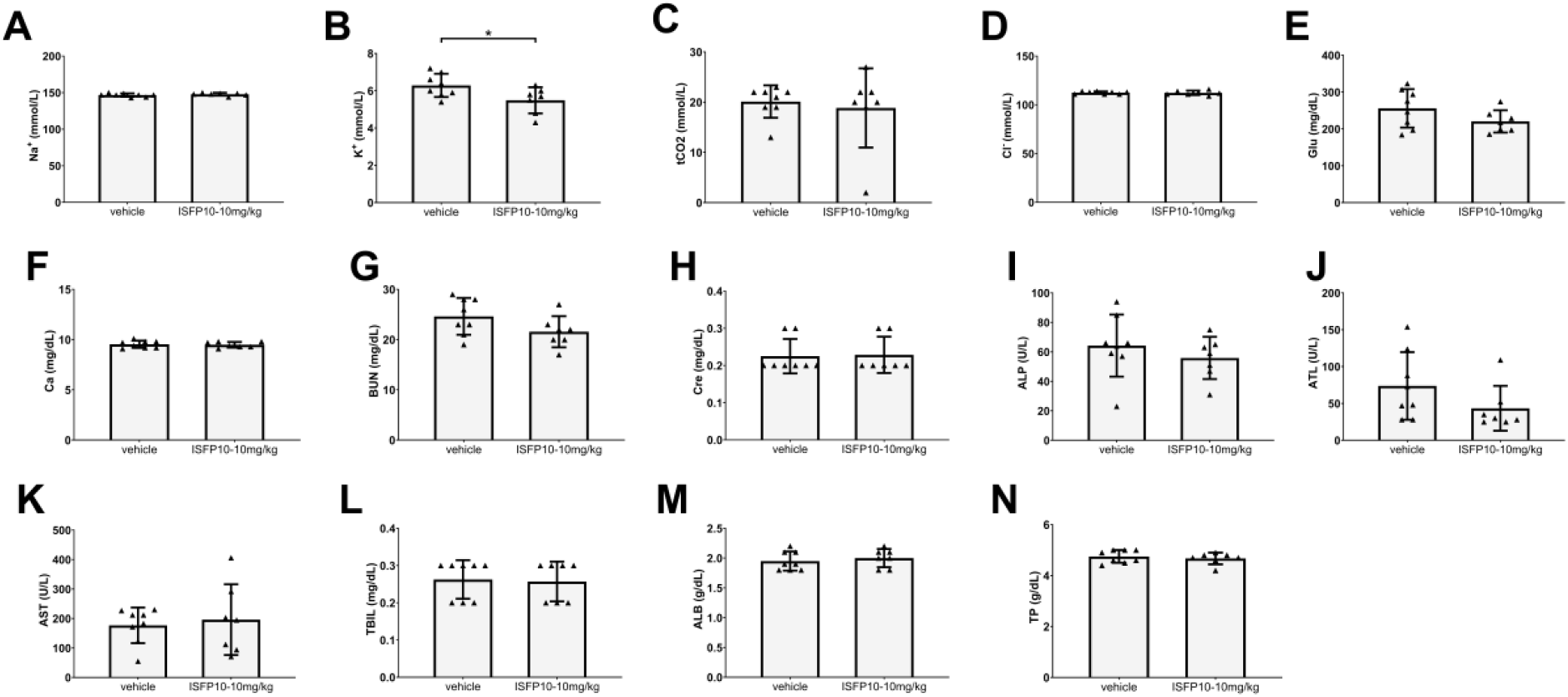
CBC analysis. WBC, total white blood cells (109/L); LYM, lymphocytes (109/L); MON, monocytes (109/L); NEU, neutrophils (109/L); RBC, red blood cells (1012/L); HGB, hemoglobin (g/dL); HCT, hematocrit (%); MCV, mean corpuscular volume (fL); MCH, mean corpuscular hemoglobin (pg); MCHC, mean corpuscular hemoglobin concentration (g/dL); PLT, platelets (109/L). (n = 7-8 mice/group). ± SD; *p < 0.05.

**Fig 5.**
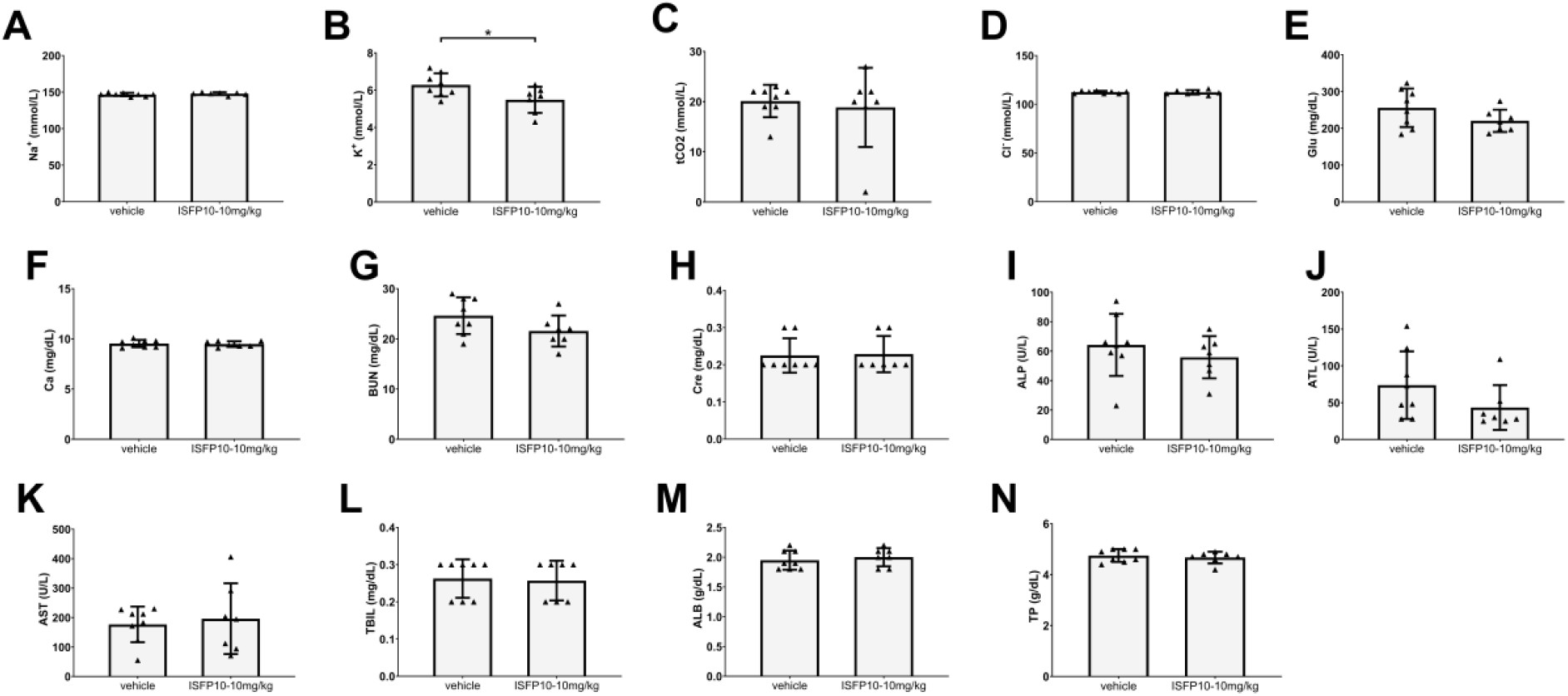
Serum chemistry parameters. Na, sodium (mmol/L); K, potassium (mmol/L); tCO2, carbon dioxide (mmol/L); Cl, chloride (mmol/L); GLU, glucose (mg/dL); BUN, blood urea nitrogen (mg/dL); Cre, creatine (mg/dL); ALP, alkaline phosphatase (U/L); ALT, alanine aminotransferase (U/L); AST, aspartate aminotransferase (U/L); TBIL, bilirubin (MG/DL); Alb, albumin (g/dL); TP, total protein (g/dL). (n = 6-8 mice/group). ± SD; *p < 0.05.

### 3.5. Histology Analysis

Histologic examination of all samples from lung, liver, and kidney from both ISFP10 treated- and vehicle groups did not reveal any tissue injury or toxic abnormalities, and all organs displayed completely normal architecture (**Fig. 6**) (score 0 for all parameters). These observations further support good tolerability of ISFP10 in the test animals.

**Fig 6.**
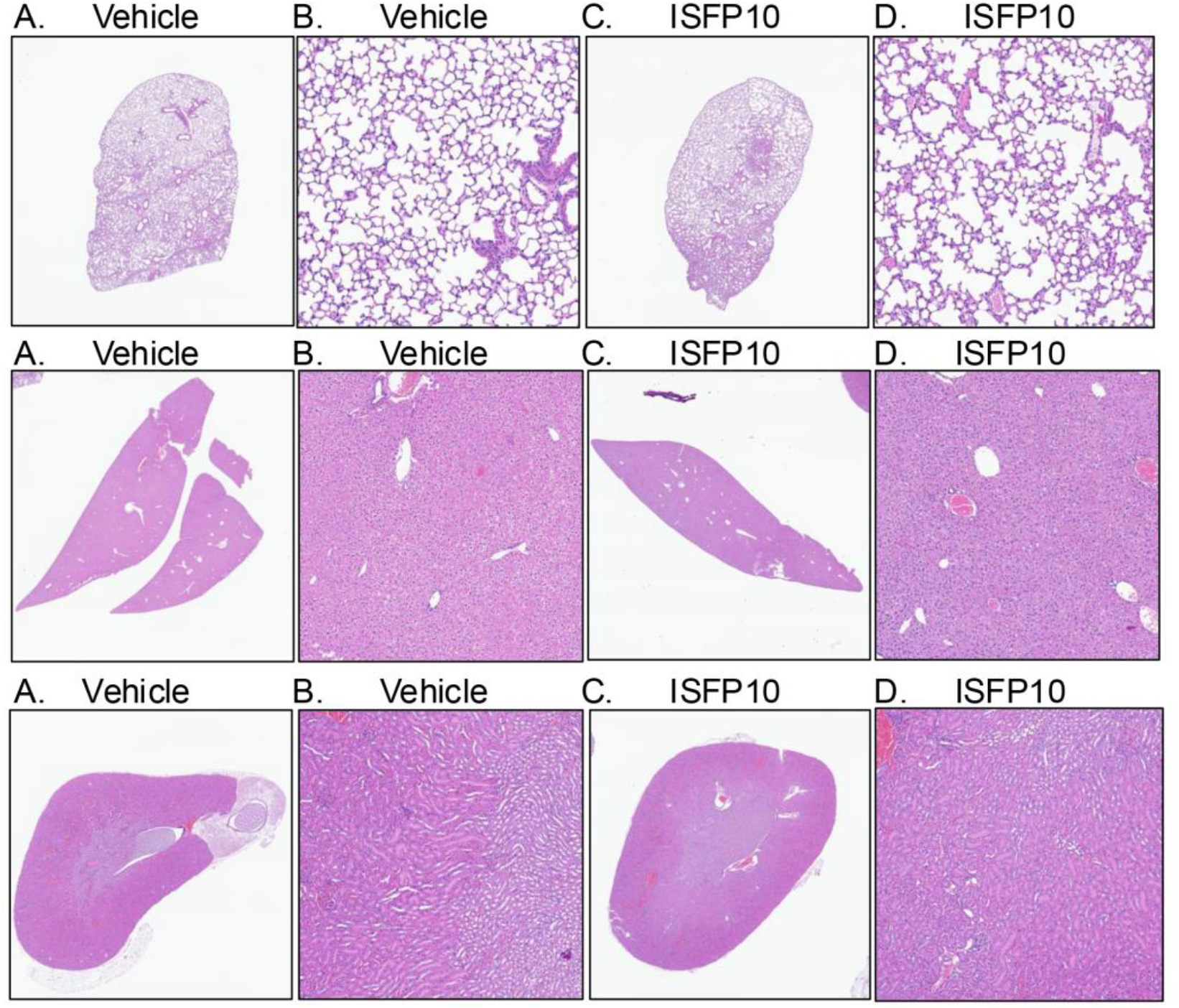
Lung, liver, and kidney histopathology of 7-day IP treated vehicle or ISFP10. Hematoxylin and eosin (H&E) staining was performed on sections of lung (top panels), liver (middle panels), and kidney (bottom panels) from mice in all groups. (A-B) Vehicle control. (C-D) ISFP10 at 10.0 mg/kg. A,C= 1X magnification. B,D= 10X magnification. No gross histological changes were present in lung, liver, and kidneys in the vehicle or ISFP10 inhibitor administration.

**Fig 7.**
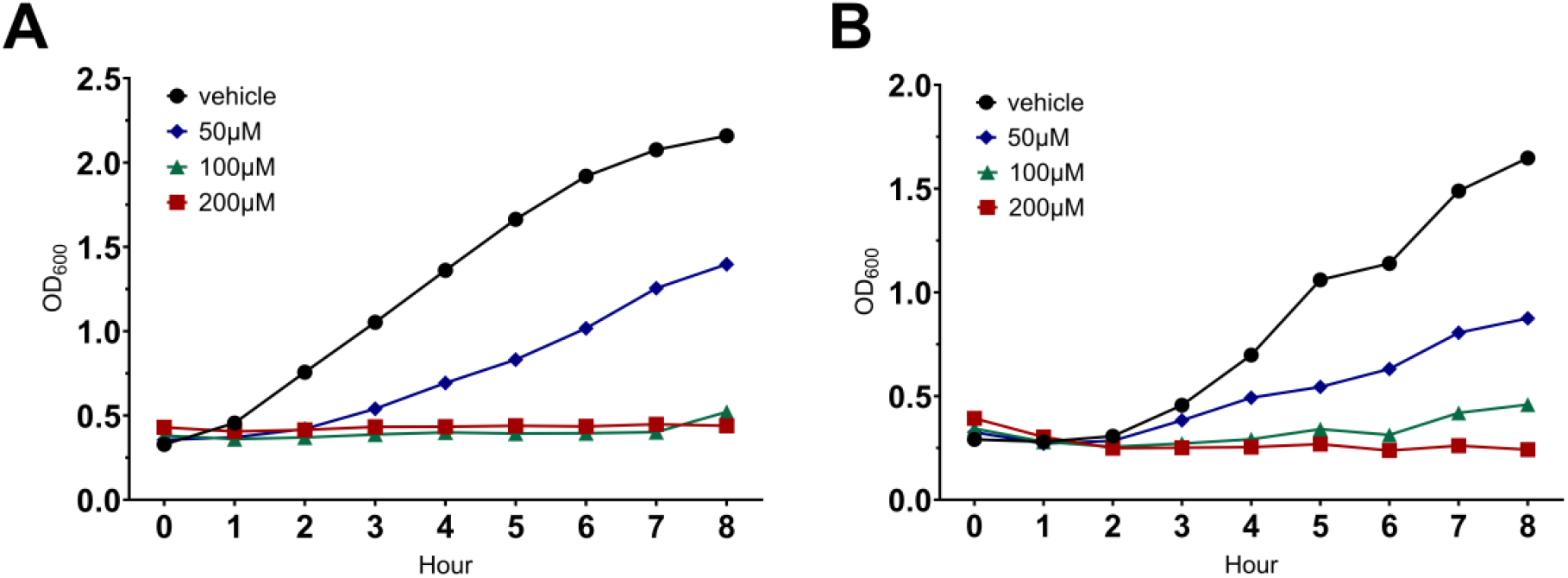
Relative growth of (A) *C. albicans* and (B) *C. glabrata*. Fungal cell growth was initiated in YPD media for 18 h at 30^°^C. After 18 h, yeast cells were diluted to an OD600_nm_ of ∼ 0.3 and ISFP10 at the noted concentrations or vehicle control added. Cells were grown for 8 h with OD600_nm_ measurements read every h for 8 h. Results shown are the mean of 3-4 independent experiments.

### 3.6. In vitro Growth Response Assay Analysis

The addition of ISFP10 to *C. albicans* and *C. glabrata* growth in YPD media versus vehicle control over 8 hours resulted in concentration-dependent inhibition of the growth of both fungal organisms (**Fig. 6** C-D). These observations add to the prior studies showing activity of ISFP10 again *A. fumigatus* and *Pneumocystis* fungal organisms, under scoring the potential broader spectrum antifungal activity of agents that inhibit fungal PGM [2, 3].

## 4. Discussion

*Candida* species including both *C. albicans* and non albicans species, colonize cutaneous and mucosal surfaces of critically ill patients [9]. Following breakdown of mucosal and immunologic defense mechanisms, systemic invasion of *Candida* can occur with serious consequences [10]. Such invasion may result in deep-seated invasive candidiasis or candidemia. *Candida* bloodstream infections can seed metastatic deposits of infections that include endovascular infections, hepatosplenic abscesses, endophthalmitis, and bone or joint infections [11, 12]. The incidence of candidemia varies in critically ill patient populations ranging from 3.5 to 16.5 episodes per 1000 ICU admissions [13-18]. Overall invasive candidiasis is associated with a high-rate mortality of 40-55% [13-15, 19]. Furthermore, concerns of emerging resistance of *Candida* species to azoles and echinocandins therapy mandates the development of new antifungal agents to treat these serious infections [20, 21].

The phosphoglucomutase (PGM) enzyme represents a potential novel and broad target for antifungal therapy. Others have shown that through the inhibition of PGM in *A. fumigatus* using the generation of a conditional *Af*PGM mutant, that the enzyme is essential for cell wall integrity and organism viability [22]. These researchers also reported via a fragment screen of 1000 compounds and scaffold exploration, that the synthesis of a para-aryl derivative PGM inhibitor, termed ISFP10, displayed an IC_50_ of 2 μM and over 50-fold selectivity over the human PGM homolog [22]. Selectivity was designed and achieved via targeting a unique cysteine residue found not only in *A. fumigatus* but also several other pathogenic fungi including *Candida* spp. and *Pneumocystis* spp. [2, 23]. ISFP10 covalently modifies and inactivates that critical cysteine in the active site of fungal PGM molecules [3]. We have further shown that ISFP10 can indeed inhibit *Pneumocystis* PGM activity, therefore making it also a possible attractive anti-*Pneumocystis* therapeutic target in addition to other pathogenic fungi [2].

To move toward clinical applications in humans and to further determine whether PGM inhibition might be a viable antifungal strategy, potentially toxic effects from the preclinical inhibitor ISFP10 must be excluded through preclinical assessment. As far as we are aware, this is the first comprehensive study conducted to investigate toxic effects of ISFP10 in test animals. Reassuringly, the data presented here demonstrates that ISFP10, is well-tolerated and remains a promising compound for further studies in animal models of fungal infections.

In this study, mice were dosed with ISFP10 at 10 mg/kg per day for seven consecutive days. The treated animals appeared to tolerate the daily IP injections well. Body weight measurements, lung ECM generation analysis, blood chemistry and CBC panel screening, assessments of lung proinflammatory cytokine release, and histological H&E examinations showed little to no clinically significant deviation from the control group. Furthermore, since many fungal PGMs contain a highly conserved active site with a targetable conserved cysteine residue, the PGM drug target may hold potential to treating a number of serious fungal infections. This was further supported by our observation that ISFP10 inhibited the growth of *C. albicans* and *C. glabrata* in broth culture over time and in a concentration dependent manner. Together, these results further support the development of ISFP10 as antifungal agent and the potential use of the fungal PGM as a therapeutic target.

## 5. Conclusions

These observations support the targeting of fungal PGMs with ISFP10 using *in vivo* models of infections, appears to be safe and merits additional preclinical development in the treatment of serious fungal infections.

## Ethics approval

This study was approved by the Mayo Clinic Institutional Animal Care and Use Committee (IACUC) approved protocol A00007622-24 on 4/1/2023.

## Consent to participate

Not applicable

## Consent for publication

Not applicable

## Availability of data and material

The datasets generated and/or analyzed during the current study are available from the corresponding author on reasonable request.

## Code availability

Not applicable

## Author contributions

M.R.P., M.B., T.J.K., S.A., E.S.Y., and A.H.L. made substantial contributions to the conception or design of the work, or the acquisition, analysis, or interpretation of data. M.R.P. and M.B. are co-first authors. All authors approved the version to be published.

